# Protein Stability Changes upon Point Mutations Identified with a Gaussian Network Model Simulating Protein Unfolding Behavior

**DOI:** 10.1101/2022.02.17.480818

**Authors:** Sambit K. Mishra

## Abstract

Understanding the effects of missense mutations on protein stability is a widely acknowledged significant biological problem. Genomic missense mutations may alter one or more amino acids leading to increased or decreased stability of the encoded proteins. In this study, we propose a novel approach - Protein Stability Prediction with a Gaussian Network Model (PSP-GNM) to study the effect of single amino acid substitutions on protein stability. Specifically, PSP-GNM employs a coarse-grained Gaussian Network Model (GNM) that has interactions between amino acids weighted by the Miyazawa-Jernigan (MJ) statistical potential. We use PSP-GNM to simulate partial unfolding of the wildtype and mutant structures and then, use the difference in energies of the unfolded wildtype and mutant protein structures to estimate the experimentally obtained unfolding free energy change (ΔΔG). We verify the extent of correspondence between the ΔΔG calculated by PSP-GNM and the ΔΔG obtained experimentally using three datasets: 350 forward mutations from 66 proteins, 2298 forward mutations from 126 proteins and 611 forward and reverse mutations from 66 proteins and observe Pearson correlation coefficient (PCC) as high as 0.58 and root mean-squared error (RMSE) as low as 1.24 kcal/mol. The performance is comparable to the existing state of the art methods. Importantly, we do observe an increase in the correlation to 0.73 and decrease in RMSE to 1.07 when considering only those measurements made close to 25°C and neutral pH, suggesting a strong dependence on temperature and pH. PSP-GNM is written in Python and is available as a free downloadable package at https://github.com/sambitmishra0628/PSP-GNM.

**Author Summary:** Understanding how genomic missense mutations impact the thermodynamic stability of encoded proteins is important to understand disease etiology. Specifically, mutant proteins are often functionally inactive and underlie numerous genetic and neurodegenerative diseases. A classic example is sickle cell anemia – a single amino acid change significantly affects hemoglobin’s binding affinity for oxygen. To be able to identify mutations that would likely impact the protein function and stability is therefore essential and is the focus of our study. We present an approach that relies on utilizing the intrinsic dynamics of protein structures to predict the effect of single amino acid mutations (point mutations) on protein stability. In our approach, we model proteins as coarse-grained beads (amino acids) and springs (interactions), simulate protein unfolding and identify putative residue-residue contacts that are broken during the unfolding process. We demonstrate that the knowledge of broken contacts and their order is essential in describing the thermodynamic differences between wildtype and mutant proteins. We also highlight the importance of residue-residue interactions at the mutation site in the context of protein stability prediction. Our findings present novel avenues to interpret how genomic mutations may manifest in the encoded proteins.

## Introduction

Genomic mutations are often random and can either be synonymous mutations that do not alter the amino acid sequence of an encoded protein or non-synonymous mutations that alter the amino acid sequence. Non-synonymous point mutations can lead to structural changes, affecting the folding energy landscape and thermodynamic stability of proteins. They may sometimes have a neutral effect on the folding energy landscape of proteins, that is there may be little difference in the thermodynamic stability of the mutant and wildtype proteins. However, mutations often tend to change the folding energy landscape rendering the mutant proteins either more or less stable compared to the wildtype. Understanding the impact of non-synonymous point mutations on the thermodynamic stability of proteins is regarded as a key biological problem. Many neurodegenerative diseases and genetic disorders have been linked to the incorrect folding of polypeptides that is caused by mutations in genes [1]. It is therefore necessary to understand how non-synonymous mutations impact the thermodynamic stability of proteins to better associate their roles in different diseases.

Mutagenesis experiments that assess the change in thermodynamic stability of proteins measure the energies of the folded and unfolded states of the wildtype and mutant proteins, providing an accurate picture of stability change upon mutations. However, such experiments are time-consuming and expensive and thus, there is a need for alternative computational methods that accurately predict the effects of mutations on protein stability. Databases, such as ProTherm [2], have mined such experimental data from the literature and have recorded details of changes in free energies (ΔΔG) upon mutations. The calculated ΔΔG is typically the difference in the Gibbs free energies (ΔG) of the mutant and wildtype proteins. Specifically, ΔΔ*G* = Δ*G*_mut_ − Δ*G*_wt_, where Δ*G*_mut_ is the energy of the unfolded state minus the energy of the folded state of the mutant protein 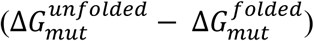, while Δ*G* is the energy of the unfolded state minus the energy of the folded state of the wildtype protein 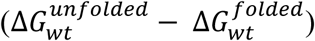 [3]. While a positive ΔΔG implies that the folded mutant protein is thermodynamically more stable than the wildtype, a negative ΔΔG indicates that the folded mutant protein is a thermodynamically less stable.

Numerous computational methods have been developed to predict the ΔΔG between a wildtype protein and its non-synonymous mutant form. The review article by Marabotti *et al*. [4] provides a comprehensive overview of the computational methods developed in this effort to quantify ΔΔG changes in proteins upon non-synonymous mutations. Existing methods may be broadly classified into two categories: 1. Unsupervised methods and 2. Supervised methods. Unsupervised methods are untrained methods and do not rely on machine learning. They can include empirical methods that model the energy of a protein through an equation that includes several parameters, each parameter carrying a different weight. For example, the FoldX empirical method [5] uses both the bonded and non-bonded energy terms to model a protein’s energy and then calculates the difference in the energies of the mutant and wildtype proteins to estimate the ΔΔG. SDM [6] uses environment-specific amino acid substitution tables to account for the amino acid substitution in the mutant, residue packing density and depth to calculate the ΔΔG. DDGun is another unsupervised method that uses sequence and structural properties of proteins in linear weighted combinations to estimate the ΔΔG [7]. Supervised methods employ machine learning to learn from a variety of features (e.g., amino acid physicochemical properties, protein geometric properties, etc.) of the wildtype and mutant forms of different proteins and associate these features with the experimentally calculated ΔΔG. Such methods are then tested and evaluated on an independent dataset of wildtype and mutant proteins with known ΔΔG.

Numerous supervised computational methods to predict stability changes in proteins upon mutations have been developed in the past decade. These methods often employ machine learning models trained on diverse benchmark datasets obtained from the Protherm database. I-mutant [8,9] employs support vector machines (SVM) [10] trained on a subset of data taken from Protherm and uses a set of sequence and spatial features, including solvent accessibility, to predict the ΔΔG. It also gives the user an option to make predictions using just the protein sequence in case the protein structure is not available. mCSM [11] represents the environment of the mutation site as a distance-based graphical map and combines it with pharmacophore counts to predict the effect of mutations using a trained linear regression model. MAESTRO [12] uses statistical scoring functions and other sequence and structural features, such as the accessible surface area, the hydrophobicity, and the secondary structure of the mutation site, to train artificial neural networks, SVM and multiple linear regression models and makes consensus ensemble-based predictions. Dynamut [13] follows a meta-prediction approach by using predictions from other methods, which use protein structure and dynamics information to estimate ΔΔG, to arrive at a consensus prediction for ΔΔG. Dynamut2 [14] predicts ΔΔG using a Random Forest algorithm trained on protein dynamics features calculated from normal mode analysis and graph-based signatures.

It has generally been observed that the supervised machine-learned methods outperform the unsupervised methods in predicting ΔΔG [14]. However, one of the key shortcomings of the supervised methods is overfitting the training data. Besides, supervised methods use an ensemble of features that are selected purely based on the algorithms’ performance in cross-validation, not based on any theoretical model that simulates the mutational perturbation behavior. It is always a challenge to associate such features with mutational perturbation, understand how a mutation affects each of these features and explain the biological significance and role of each feature in the context of protein thermodynamic stability.

In this study, we present a theoretical model - PSP-GNM, which simulates protein unfolding behavior using the Gaussian Network Model (GNM) [15], to estimate ΔΔG upon non-synonymous mutations. GNM is an elastic network model that models a protein using a coarse-grained representation. Each amino acid in GNM is represented by its alpha carbon and the interacting amino acid pairs are connected using hypothetical Hookean springs [15]. Previously, elastic network models have been shown to capture the near-native dynamics of proteins [16] and have been used to efficiently identify functional sites in proteins [17,18]. GNM has been previously employed to study changes in protein dynamic communities upon mutations and has been shown to possess the ability to identify stabilizing and destabilizing mutations in proteins [19]. In addition, GNM has also been previously utilized to study the protein unfolding behavior and identify the order of contacts broken during protein unfolding [20,21]. PSP-GNM utilizes the knowledge of amino acid contacts that are broken at the mutation site during theoretical protein unfolding to estimate the energy change. It is based on protein dynamics obtained using a coarse-grained representation of the protein and provides useful insights into the importance of the local dynamics of the mutation site and its interactions with the neighbors to understand protein stability changes upon mutation. We have evaluated the calculated ΔΔG from PSP-GNM on multiple benchmark datasets obtained from the Protherm database and compared the predictions from PSP-GNM with other state of the art methods. Despite being an unsupervised approach, PSP-GNM shows comparable performance to many supervised approaches.

## Materials and Methods

### Datasets

We used three benchmark datasets that were originally created from the ProTherm database and have been widely used by other methods. Specifically, these datasets were procured from Dynamut2 [14], a recently developed supervised prediction method that uses them to test the performance of multiple methods. Each dataset includes non-synonymous point mutations and the experimentally measured ΔΔG for a set of proteins with experimentally resolved protein structures.

**1**. The S2298 dataset comprises 2298 forward mutations across 126 unique PDB IDs. It is originally a subset of the S2648 dataset that comprises 2648 non-synonymous point mutations across 131 unique proteins and has been widely used by supervised methods to train prediction algorithms. **2**. The original S350 dataset comprises 350 forward mutations across 67 unique PDB IDs. It is a subset of the S2648 dataset and has been frequently used for testing by other methods. We excluded mutations for PDB ID 2A01 as the entry for 2A01 was obsolete in PDB. Consequently, we had 349 mutations from 66 PDB IDs. **3**. The S611 dataset originally includes 611 forward and reverse mutations across 67 unique PDB IDs. Upon exclusion of mutations for the PDB ID 2A01, we have 609 mutations from 66 unique PDB IDs. Details of the proteins and mutations in each of the datasets is included in the supporting documents (S350.csv, S611.csv and S2298.csv).

A larger fraction of mutations in each dataset are destabilizing mutations i.e., ΔΔG < 0 kcal/mol. The S2298 dataset comprises 1791 destabilizing mutations, 478 stabilizing mutations (ΔΔG > 0 kcal/mol) and 29 neutral mutations (ΔΔG = 0 kcal/mol). The S350 dataset comprises 259 destabilizing mutations, 85 stabilizing mutations and five neutral mutations. While both the S2298 and S350 dataset include only forward mutations, the S611 dataset includes both the forward and reverse mutations. In this context, if X-N-Y denotes a forward mutation from amino acid X to amino acid Y at position N and has ΔΔG = k, then the reverse mutation (Y-N-X) is assumed to be anti-symmetric to the forward mutation and has ΔΔG = -k. The S611 dataset is specifically used to evaluate the extent of agreement with this symmetric association of forward and reverse mutations for different methods. It includes 330 destabilizing mutations, 269 stabilizing mutations and 10 neutral mutations.

### PSP-GNM algorithm

We represented the wildtype and mutant protein structures as coarse-grained systems where each residue was represented by its alpha carbon. Additionally, we represented the mutant protein structures by replacing the wildtype residues with their respective mutant residues at the alpha carbon position. We then took a similar approach using GNM as previously described by Su *et al*. [20] to simulate protein unfolding and identify the sequence of residue-residue contacts broken during unfolding. Specifically, Su *et al*. had demonstrated protein unfolding using GNM for two proteins (Barnase and Chymotrypsin inhibitor) and observed significant agreement in the sequence of unfolding events with atomic molecular dynamics and Monte Carlo simulations. Like their approach, we have also used a weighted-GNM where the strength of springs between interacting residues varies by the type of residue pair

In GNM, the protein residues are represented by alpha carbons and residue pairs within a cutoff distance r_c_ are hypothetically connected with Hookean springs of force constant γ=1. In PSP-GNM we weight the interactions between residues by their corresponding Miyazawa-Jernigan (MJ) contact potential [22], which was obtained from the AAindex database [23] using the identifier MIYS960101. Specifically, the contact potential for a residue pair is converted into its Boltzmann weight (by simply taking its exponential) and is then used to weight the interaction between the residue pair. The potential for PSP-GNM is given as

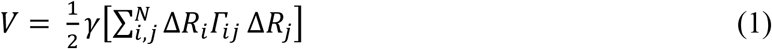

where Δ*R*_*i*_ is the fluctuation vector for residue *i* and Δ*R*_*j*_ is the fluctuation vector for residue *j*. The force constant *γ* is weighted for each interacting residue pair depending on the type of interaction. *Γ*_*ij*_ is the *ij*th element in the Kirchhoff matrix of inter-residue contacts and is given as follows.

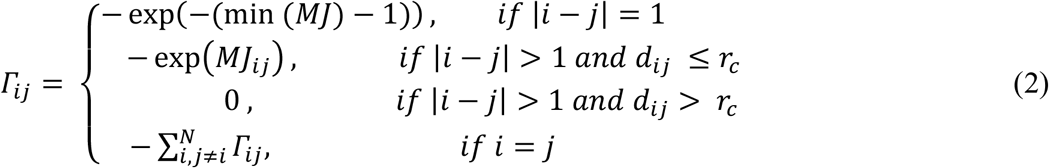

In equation 2 *MJ*_*ij*_ is the contact energy between residue *i* and residue *j* as given in the MJ potential matrix, *d*_*ij*_ is the distance between the alpha carbons of residues *i* and *j* and *r*_*c*_ is the distance cutoff for interacting residues. Stiffer springs were used to simulate the stronger covalent interactions (|*i* − *j*| = 1) between adjacent residue pairs by using the minimum value in the MJ potential matrix minus one.

To simulate unfolding, we iteratively identify residue-residue contacts with the highest mean-squared fluctuations (MSF) in their distance [20,24] and then remove the contacts between these residues. The MSF in distance between residue pairs *i* and *j* is calculated as

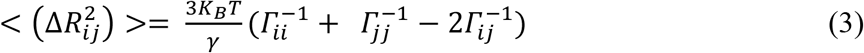

where, *K*_*B*_ is the Boltzmann constant and T is the absolute temperature in Kelvin. *Γ*^-1^ is the pseudo-inverse of the Kirchhoff matrix and is given as

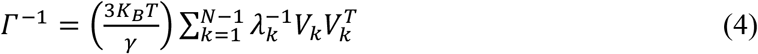

As the determinant of the Kirchhoff matrix is zero, its inverse cannot be directly calculated. Instead, we calculate the pseudo-inverse using the *N* – 1 non-zero eigen values and their corresponding eigen vectors. In the above equation, *λ*_*k*_ is the *k*th eigen value, *V*_*k*_ is the *k*th eigen vector associated with *λ*_*k*_ and 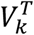 its transpose.

The steps in our algorithm are outlined in Fig. 1. Given the wildtype or mutant C-alpha PDB structure, we first identify the total number of residue-residue contacts in the starting structure. A contact pair is a residue pair whose C-alpha atoms are within 9Å. We then calculate the MSF in distance between all residue pairs by using only the 10 slowest non-zero modes to re-construct the *Γ*^-1^. The above values for the r_c_ (C-alpha distance cutoff) and number of modes parameters were used as they resulted in maximum agreement between the experimental ΔΔG and PSP-GNM-calculated ΔΔG. The contact pair having the largest MSF in distance is then identified, the contact between the residue pair broken and the contact matrix is updated to reflect this contact break. The updated contact matrix is then used in the next iteration to identify the contact pair with the largest MSF and highest likelihood of contact break. We repeat this scheme of breaking contacts based on the largest MSF until 50% of all contacts in the starting structure are broken, resembling a partially unfolded protein, or until the model becomes unstable resulting in more than one 0-eigen values upon eigen decomposition of the Kirchhoff matrix.

**Figure 1.**
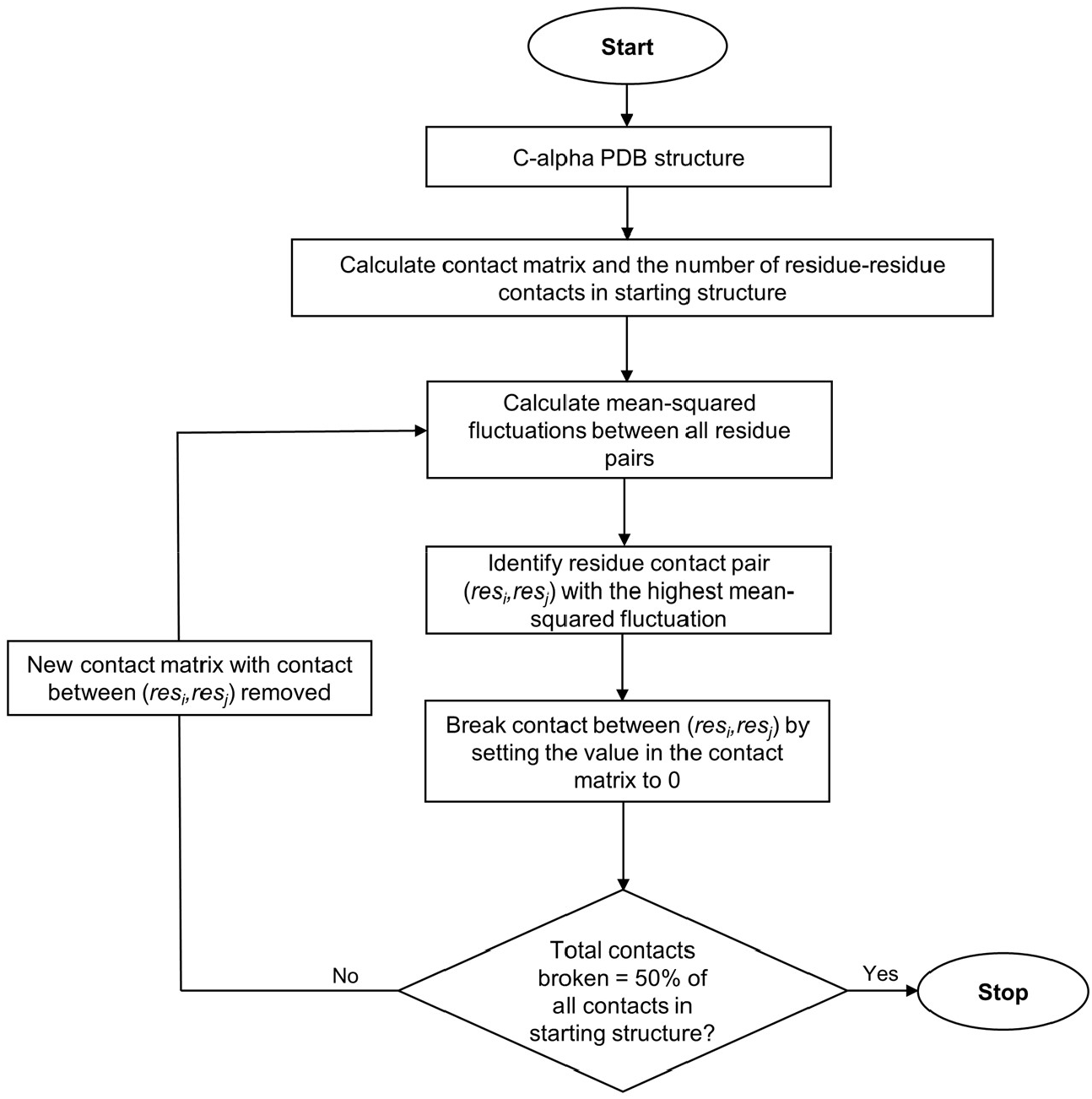
Overview of PSP-GNM. .The steps involved in PSP-GNM are outlined in this figure. The input is a coarse-grained protein structure that includes only the residue C-alpha atoms. The protein is then partially unfolded by sequentially identifying residue contact pairs with the highest mean-squared fluctuations in their distance and removing the contacts between these residue pairs. The process is repeated until 50% of native contacts are broken.

### Calculation of ΔΔG using PSP-GNM

PSP-GNM assumes a given mutant protein differs from its corresponding wildtype by only one amino acid. That is, the current implementation of PSP-GNM can be used to calculate ΔΔG given a mutant structure with a single amino acid change compared to its wildtype. For a pair of wildtype and mutant proteins, the contacts broken with the residue at the mutation position during partial unfolding are serially ranked (starting with a rank of 1) in the order of which they were broken. The underlying assumption is that there is at least a single contact involving the residue at mutation position that is broken during the partial unfolding of the mutant and wildtype structures. Then, we only consider the set of broken contacts in the mutant and wildtype that share the same rank. ΔΔG is then calculated as

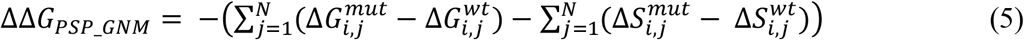

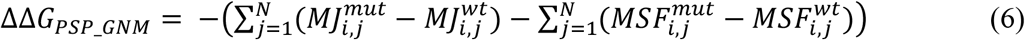

In the above, 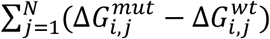 is the total difference in interaction energies of the mutant and wildtype, 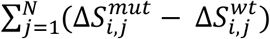 is the difference in entropy. 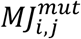 and 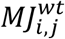 are the Miyazawa-Jernigan interaction energies between residue at mutation position *i* and the residue with which contact was broken having rank *j* in the mutant and wildtype proteins, respectively.

The MSF in distance is considered as a measure of entropy. 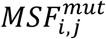 and 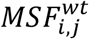 are the mean-squared fluctuations between residue at mutation position *i* and the residue with which contact was broken having rank *j* in the mutant and wildtype proteins, respectively. Re-arranging the terms, Eq. 5 can be re-written as

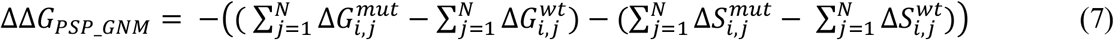

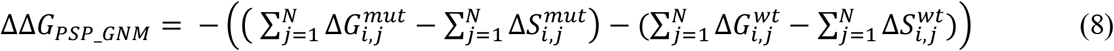

During partial unfolding, if no contacts involving the residue at the mutation position are broken either in the wildtype structure or in the mutant structure or both, then ΔΔG cannot be calculated. For such cases we assign a theoretical ΔΔG value of 0. The calculated ΔΔG are fit to the experimental ΔΔG using linear regression and the equation of the line of best fit is used to re-scale the calculated ΔΔG.

### Calculation of ΔΔG for reverse mutations

To perform calculations for reverse mutations (ΔΔG_*mutant*_ → ΔΔG_*wildtype*_), we simply switched the wildtype and the mutant structures. Then, we followed the same steps as described above to calculate ΔΔG.

### Evaluation criteria

To estimate the extent of agreement between the ΔΔ*G*_PSP_GNM_ with the experimental ΔΔG we used Pearson correlation coefficient and root mean squared error. Pearson correlation was calculated using the scipy.stats module in Python (https://docs.scipy.org/doc/scipy-0.14.0/reference/generated/scipy.stats.pearsonr.html#scipy.stats.pearsonr), while RMSE was calculated using the sklearn.metrics.mean_squared_error module in Python (https://scikit-learn.org/stable/modules/generated/sklearn.metrics.mean_squared_error.html).

## Results

PSP-GNM utilizes coarse-grained Gaussian Network Model (GNM) to simulate partial protein unfolding. Unlike the conventional GNM that has interacting residues connected with springs having a uniform force constant of 1, PSP-GNM weights the interacting residues by their interaction energies obtained from the Miyazawa-Jernigan potential [22]. To simulate partial protein unfolding, residue-residue contacts with the highest MSF in their distance are iteratively identified and broken. We rank the contacts broken with the mutation site during partial unfolding in the order of which they were broken with respect to the mutation site. Then we only consider the broken contacts that share the same rank in the mutant and wildtype protein for calculations of ΔΔG. A key underlying assumption is that the MSF in distance between residue pairs is a measure of their entropy. The difference between the energy and entropy terms of the mutant and wildtype proteins constitutes the calculated ΔΔG by PSP-GNM. Details are included in Materials and Methods.

First, we identify the distance and number of modes parameters for which PSP-GNM shows maximum agreement with experimental ΔΔG. Additionally, we investigate the extent of agreement between the residue mean-squared fluctuations calculated by our method with the experimental temperature factors. Second, we evaluate the extent of agreement of PSP-GNM-calculated ΔΔG with experimental ΔΔG on different datasets. Finally, we compare the performance of PSP-GNM with other state of the art methods.

### Parameters for PSP-GNM

Two essential parameters regulating the protein dynamics captured by PSP-GNM are the cutoff distance (r_c_) between C-alpha atoms and the number of low-frequency modes (N_modes_) used to reconstruct the inverse Kirchhoff matrix. While r_c_ defines the number of interacting residue-pairs in a protein, N_modes_ impacts the calculated MSF in distance between a residue pair. We identify the values for these two parameters that maximize the agreement between the PSP-GNM-calculated ΔΔG and the experimental ΔΔG on the S350 dataset. The choice for using the S350 dataset for this purpose is arbitrary. First, we set the N_modes_ to 10 and identify the r_c_ in the range of 7Å-12Å that results in maximum correlation between PSP-GNM-calculated ΔΔG and the experimental ΔΔG. We observe the highest Pearson correlation coefficient of 0.51 for r_c_ = 9Å (Fig. 2A). Then, we set r_c_ to 9Å and observe the highest correlation of 0.51 for N_modes_=10 when using 5, 10, 20, 30, 40 and 50 modes (Fig. 2B). Therefore, we perform all our analysis using r_cutoff_ = 9Å and N_modes_=10.

**Figure 2.**
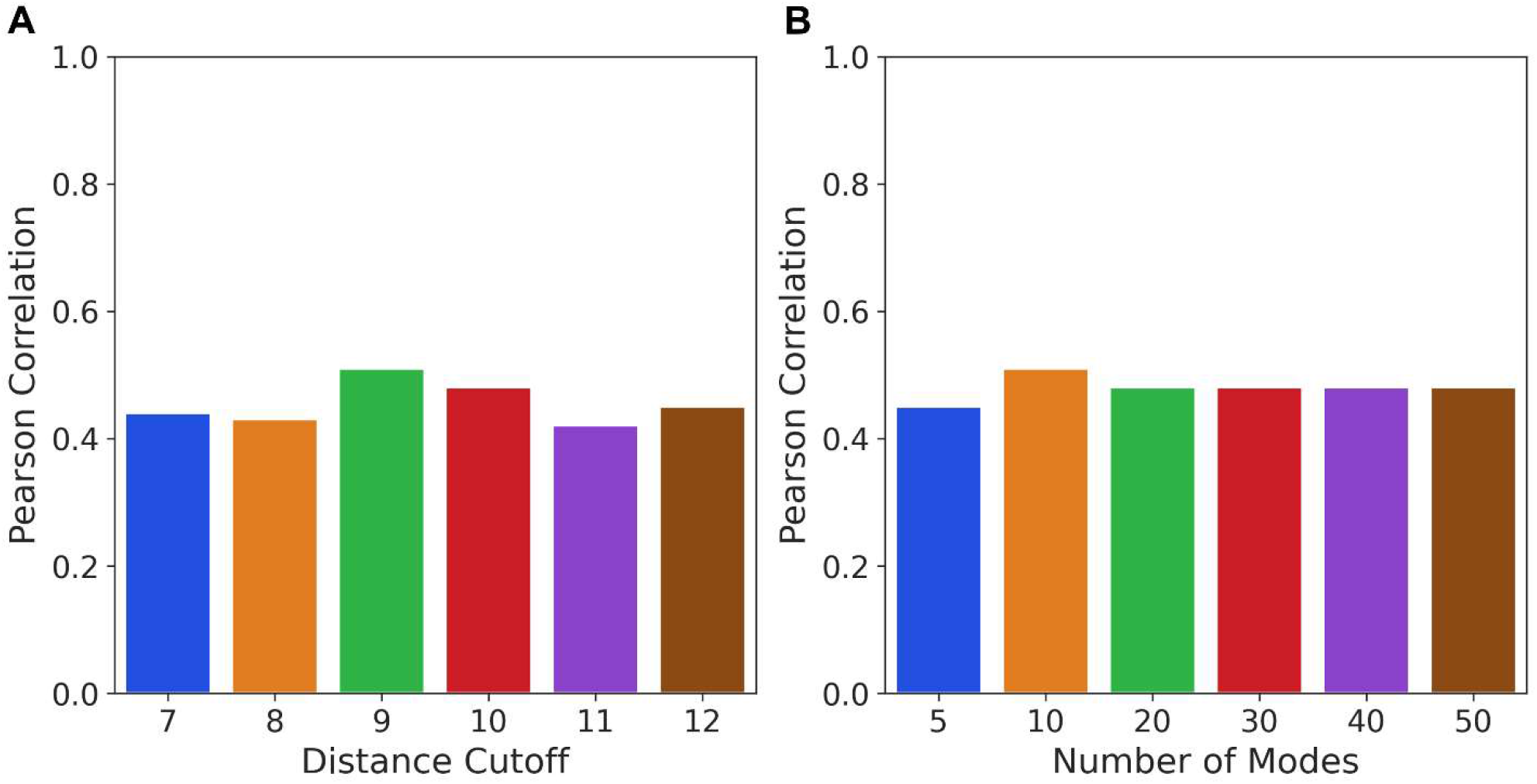
Optimal parameters for PSP-GNM. The S350 dataset is probed to identify optimal parameters for the cutoff distance for residue-residue contacts **(A)** and the total number of modes used to calculate the mean-squared fluctuation in residue-residue distance **(B)**. The agreement is best for a distance cutoff of 9Å and when using the top 10 non-zero low frequency modes.

We also investigate the extent of agreement of the residue mean-squared fluctuations obtained using PSP-GNM with the experimental B-factors. We perform this comparison on 66 wildtype proteins from the S350 dataset that are represented as C-alpha coarse-grained structures. Only 57 proteins were X-ray crystal structures and included B-factors. We observe a median correlation of 0.48 between the mean-squared fluctuations calculated by PSP-GNM and the experimental B-factors (Fig. S1). We also note that this agreement is marginally weaker than GNM (correlation 0.58), when using r_c_ = 9Å and N_modes_=10 (Fig. S1).

### Predictive performance of PSP-GNM on different datasets

We evaluated the extent of agreement for the PSP-GNM-calculated ΔΔG with the experimental ΔΔG on three datasets: 1. S2298 dataset includes 2298 forward mutations across 126 unique PDBs, S350 dataset that includes 349 mutations from 66 PDBs, and 3. S611 dataset that includes 609 forward and reverse mutations across 66 unique PDBs. In the following text, we report the performance on each dataset.

#### Performance on S2298 dataset

Of the 2298 mutants in the S2298 dataset, PSP-GNM-calculated ΔΔG could be obtained for 2155. For these 2155 mutants, there was at least one contact involving the mutation position that was broken during partial unfolding of the mutant and wildtype structures, permitting ΔΔG calculations. We observe a Pearson correlation of 0.39 and RMSE of 1.35 kcal/mol when compared to the experimental ΔΔG (Fig. 3A). Upon inclusion of the 143 mutations without any PSP-GNM-calculated ΔΔG (by assigning them a theoretical ΔΔG of 0 kcal/mol), we do not observe any considerable change in the agreement with the experimental data (Fig. 3B).

**Figure 3.**
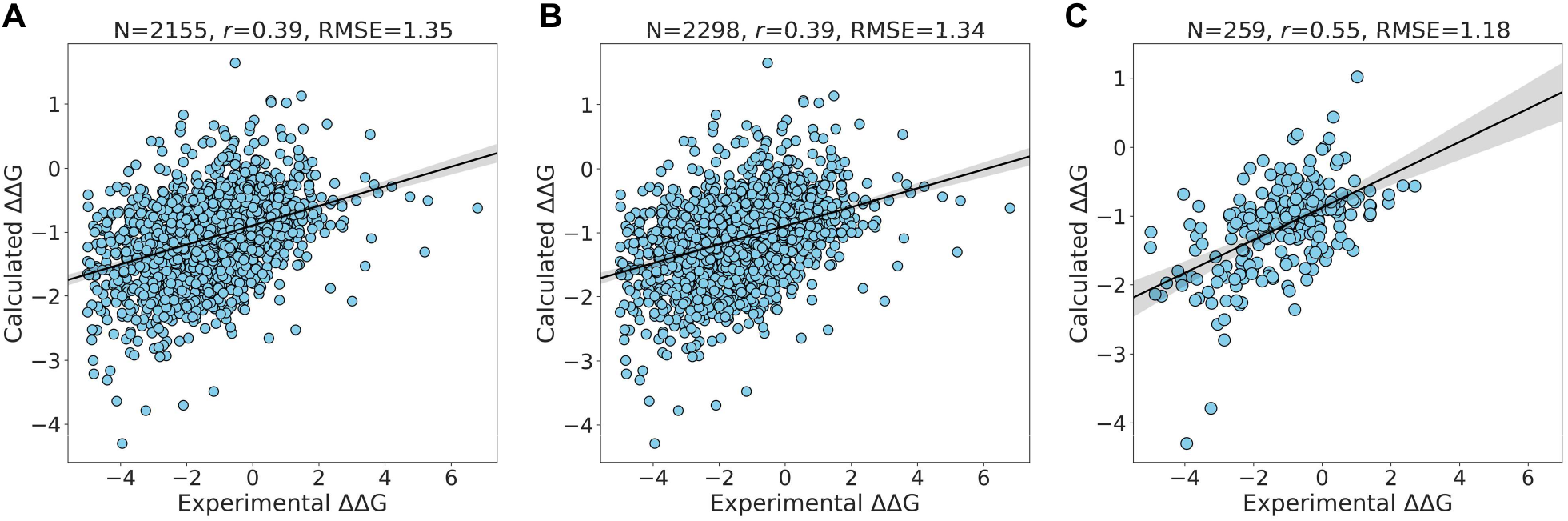
Performance on S2298 dataset. The extent of agreement between the experimental ΔΔG and the PSP-GNM-calculated ΔΔG is shown for **(A)** 2155 mutants having a PSP-GNM-calculated ΔΔG, **(B)** All 2298 mutants, **(C)** 259 mutants with experimental temperatures ranging from 24°C - 26°C and pH ranging from 6.8 - 7.2. The unit for the calculated ΔΔG is kcal/mol. The regression line with the 95% confidence interval (shaded gray) is shown for all the three cases. The optimal agreement is seen for **(C)** with a Pearson correlation of 0.55 and RMSE of 1.18 kcal/mol. We use a theoretical ΔΔG value of 0 kcal/mol for cases without a PSP-GNM-calculated ΔΔG.

The distribution of the experimental temperature and pH for the 2155 mutations with calculated ΔΔG indicates a peak around 25°C and a pH close to 7 (Fig. S2), suggesting that the experimental measurements were performed mostly around this temperature and pH. We then test whether PSP-GNM shows better agreement to ΔΔG measurements performed around 25°C and a pH of 7. Of the 2155 mutations, we identified 259 mutations with experimental temperatures ranging from 24°C - 26°C and pH ranging from 6.8 - 7.2 and observe an improved agreement (Pearson correlation = 0.55, RMSE = 1.18 kcal/mol) with the experimental measurements (Fig. 3C). Further, we evaluate the extent of agreement for measurements made between 24°C - 26°C at varying ranges of pH and measurements made close to neutral pH but having a broad range of temperature (Fig. S3). We observe that the temperature dependency for PSP-GNM is greater than that of pH dependency. PSP-GNM demonstrates stronger agreement with measurements made between 24°C - 26°C but, having a broad range of pH (Pearson correlation = 0.45, RMSE = 1.3 kcal/mol) than with measurements made at neutral pH that are associated with a broad range of temperature (Pearson correlation = 0.36, RMSE = 1.41 kcal/mol).

#### Performance on S350 dataset

The S350 dataset includes 349 mutations from 66 unique PDB identifiers. On this dataset, ΔΔG could be calculated with PSP-GNM for 318 wildtype-mutant pairs that exhibited at least one broken contact involving the mutation position during partial unfolding. We observe a Pearson correlation of 0.51 and RMSE of 1.36 kcal/mol for these 318 mutations when compared to the experimental ΔΔG (Fig. 4A). When the remaining 31 wildtype-mutant pairs are included by assigning them a ΔΔG value of 0 kcal/mol, we observe a decrease in the Pearson correlation to 0.49 while the RMSE still remains at 1.36 kcal/mol (Fig. 4B).

**Figure 4.**
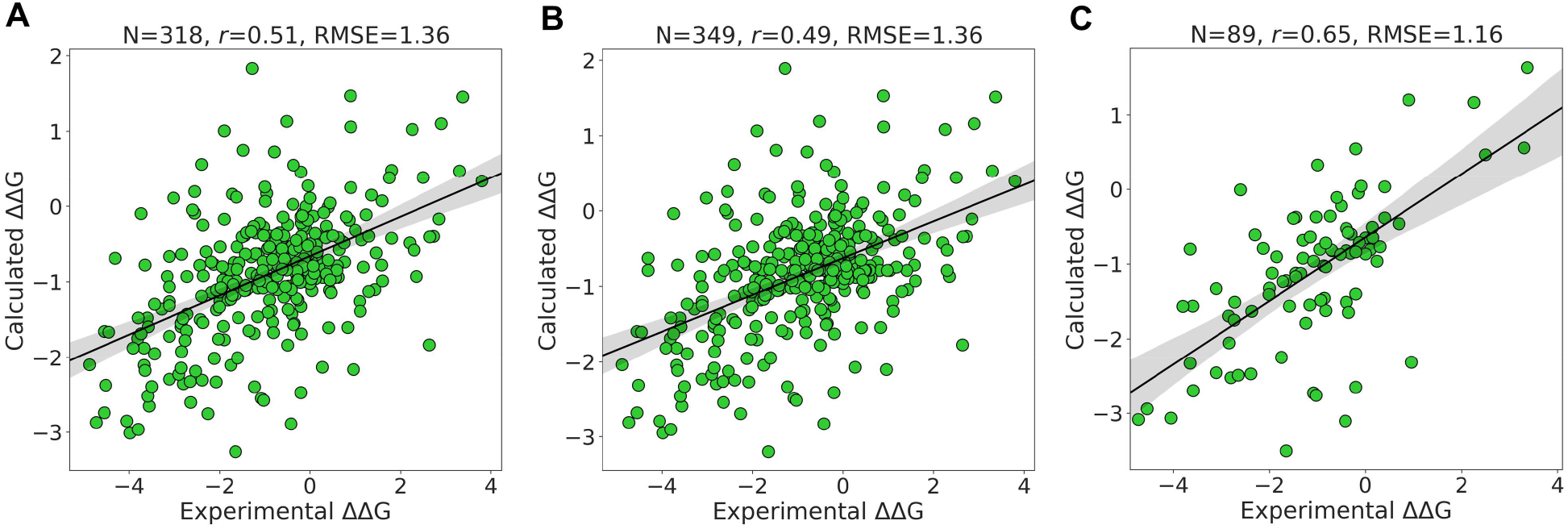
Performance on S350 dataset. Agreement between the experimental ΔΔG and the PSP-GNM-calculated ΔΔG is shown for **(A)** 318 mutants with a PSP-GNM-calculated ΔΔG, **(B)** All 349 mutants, **(C)** 89 mutants with experimental temperatures ranging from 24°C - 26°C and pH ranging from 6.8 - 7.2. The unit for the calculated ΔΔG is kcal/mol. The regression line with the 95% confidence interval (shaded gray) is shown for all the three cases. The optimal agreement is seen for **(C)** with a Pearson correlation of 0.65 and RMSE of 1.18 kcal/mol.

The experimental temperature and pH for the 318 mutants with calculated ΔΔG show a similar distribution as that of the S2298 dataset i.e., a peak around 25°C and pH 7 (Fig. S4). We observe a considerably stronger agreement with the experimental ΔΔG when considering a subset of 89 mutations with experimental temperature 24°C - 26°C and pH 6.8 - 7.2, than when considering all the data (Fig. 4C). The associated Pearson correlation and RMSE are 0.65 and 1.16 kcal/mol, respectively. The agreement is stronger when we consider a temperature range of 24°C - 26°C but no strict pH cutoff (Pearson correlation = 0.57, RMSE = 1.37 kcal/mol) than when we consider a pH range of 6.8 - 7.2 but no strict temperature cutoff (Pearson correlation = 0.25, RMSE = 1.15 kcal/mol) (Fig. S5).

Additionally, we investigate the extent to which PSP-GNM satisfies the anti-symmetric property of forward and reverse mutations using the S350 dataset. Ideally, the forward and reverse mutations should be symmetric in terms of their ΔΔG, i.e., ΔΔG_*wt→mut*_ = - ΔΔG_*mut→wt*_, where ΔΔG_*forward*_ *=* ΔΔG_*wt→mut*_ and ΔΔG_*reverse*_ = ΔΔG_*mut→wt*_. To calculate the ΔΔG_*reverse*_ we switch the wildtype and mutant structures and then simulate partial unfolding for each wildtype-mutant pair. We observe a strong negative correlation between the calculated ΔΔG_*forward*_ and ΔΔG_*reverse*_ (Pearson correlation = 0.95, Fig. S6), suggesting that the calculations by PSP-GNM satisfy the expected anti-symmetric property.

#### Performance on S611 dataset

In contrast to the S350 and S2298 datasets, the S611 dataset evaluates the performance of PSP-GNM on both forward and reverse mutations. The S611 is created using a subset of data from the S350 dataset and comprises 609 mutations from 66 unique PDB identifiers. PSP-GNM could calculate ΔΔG for 554 wildtype-mutant pairs as the remaining 55 pairs did not involve a contact break for the residue at the mutation position.

When considering the calculated and experimental ΔΔG for the 554 mutations, we observe a correlation of 0.58 and RMSE of 1.24 kcal/mol (Fig. 5A) demonstrating stronger agreement than observed for the 318 forward mutations in the S350 dataset. As seen in Fig. 5B, when we include the 55 mutations without PSP-GNM-calculated ΔΔG, we do not observe considerable changes in the agreement (Pearson correlation = 0.57, RMSE = 1.22 kcal/mol). However, when we limit the analysis to only those mutations with experimental temperatures 24°C - 26°C and pH 6.8 - 7.2, we note a considerable improvement in the correlation to 0.73 and a reduction in the RMSE to 1.07 kcal/mol (Fig. 5C). Additionally, we note that the agreement is slightly better when restricting to measurements made between 24°C - 26°C without any pH filter (Pearson correlation = 0.63) compared to measurements made near neutral pH (6.8 - 7.2) without a temperature filter (Pearson correlation = 0.61) (Fig. S7).

**Figure 5.**
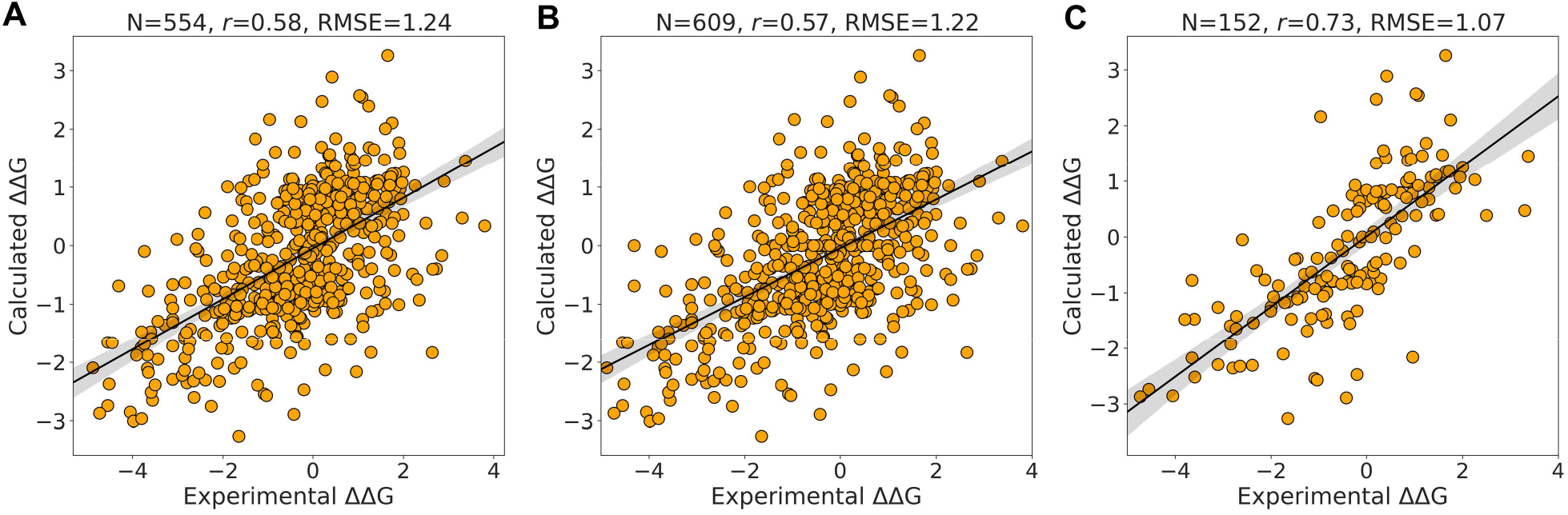
Performance on S611 dataset. Agreement between the experimental ΔΔG and the PSP-GNM-calculated ΔΔG is shown for **(A)** 554 mutants with a PSP-GNM-calculated ΔΔG, **(B)** All 609 mutants, **(C)** 152 mutants with experimental temperatures ranging from 24°C - 26°C and pH ranging from 6.8 - 7.2. The unit for the calculated ΔΔG is kcal/mol. The regression line with the 95% confidence interval (shaded gray) is shown for all the three cases. The optimal agreement is seen for **C** with a Pearson correlation of 0.73 and RMSE of 1.07 kcal/mol.

### Performance comparison with existing methods

We evaluate the performance of PSP-GNM with existing nine methods. Specifically, we assess the performance against Dynamut2 [14], Dynamut [25], MCSM [11], MUPro [26], ENCOM [27], DUET [28], SDM [6], I-Mutant2 [8], Maestro [12]. While ENCOM and SDM are unsupervised methods, the remaining methods use machine learning for predicting ΔΔG. The recently developed Dynamut2 approach also considers the above methods for comparison, and for this study we consider them as they are a good mixture of supervised and unsupervised methods. We use the S350 and S611 datasets to assess the performance of the different methods. We note that most of the existing methods have used the S2298 dataset as a training set. Consequently, a direct comparison on the S2298 dataset is not possible.

For each dataset, comparisons are made across methods by considering two groups of mutations: *Group 1*. Only those mutations that have a PSP-GNM-calculated ΔΔG, *Group 2*. Those mutations in Group 1 with an experimental temperature ranging from 24°C - 26°C and pH ranging from 6.8 - 7.2.

#### S350 dataset

In Table 1, we report the comparisons made for Group 1 across the 10 methods. Of the total 349 mutations in the S350 dataset, the comparisons are made using the 318 wildtype-mutant pairs for which there was a ΔΔG calculated. PSP-GNM ranks fifth with a correlation of 0.51 and RMSE of 1.36 kcal/mol, while DUET shows the best performance with correlation 0.67 and RMSE 1.17 kcal/mol. When we perform the same comparison on Group 2 mutants that has a total number of 89 mutants, PSP-GNM ranks fourth, showing a correlation of 0.65 and RMSE 1.17 kcal/mol, while Dynamut2 and DUET top the list with a correlation of 0.73 and RMSE 1.05 kcal/mol (Table 2).

**Table 1.** Performance on S350 dataset for a subset of 318 mutations with a PSP-GNM calculated ΔΔG.

| Method | Pearson Correlation | RMSE (kcal/mol) |
| --- | --- | --- |
| DUET | 0.67 | 1.17 |
| MCSM | 0.66 | 1.20 |
| Dynamut2 | 0.65 | 1.20 |
| IMutant | 0.54 | 1.34 |
| **PSP-GNM** | **0.51** | **1.36** |
| SDM | 0.48 | 1.91 |
| Dynamut | 0.40 | 1.79 |
| MUPro | 0.18 | 1.64 |
| ENCOM | -0.15 | 2.57 |
| Maestro | -0.57 | 1.94 |

**Table 2.** Performance on S350 dataset for a subset of 89 mutations having a calculated ΔΔG and with experimental temperatures ranging from 24°C - 26°C and pH ranging from 6.8 - 7.2.

| Method | Pearson Correlation | RMSE (kcal/mol) |
| --- | --- | --- |
| Dynamut2 | 0.73 | 1.05 |
| DUET | 0.73 | 1.05 |
| MCSM | 0.71 | 1.08 |
| **PSP-GNM** | **0.65** | **1.17** |
| SDM | 0.63 | 1.68 |
| Dynamut | 0.55 | 1.70 |
| IMutant | 0.49 | 1.33 |
| MUPro | 0.23 | 1.65 |
| ENCOM | -0.54 | 2.31 |
| Maestro | -0.68 | 2.07 |

#### S611 dataset

In Table 3, we report the comparisons for Group 1 mutants from the S611 dataset. The total number of mutations taken into consideration in this group is 554. Remarkably, we observe that PSP-GNM performs better than most of the other methods by ranking second and having a correlation of 0.58 and an RMSE of 1.24 kcal/mol. The best performance is seen for Dynamut2 (correlation = 0.68, RMSE = 1.09 kcal/mol). There were 152 mutants that were considered in Group 2, and we observe the best performance for PSP-GNM across all datasets in this group (Table 4). Specifically, PSP-GNM demonstrates strong agreement with the experimental ΔΔG by ranking second and with a correlation of 0.73 and RMSE of 1.07 kcal/mol. Overall, the best performance for this group is seen for Dynamut2 (correlation 0.79 and RMSE 0.95 kcal/mol).

**Table 3.** Performance on S611 dataset for a subset of 554 mutations with a PSP-GNM calculated ΔΔG.

| Method | Pearson Correlation | RMSE (kcal/mol) |
| --- | --- | --- |
| Dynamut2 | 0.68 | 1.09 |
| **PSP_GNM** | **0.58** | **1.24** |
| DUET | 0.50 | 1.43 |
| MCSM | 0.47 | 1.46 |
| SDM | 0.36 | 1.96 |
| IMutant | 0.34 | 1.51 |
| Dynamut | 0.27 | 1.53 |
| MUPro | 0.15 | 1.70 |
| ENCOM | -0.14 | 2.09 |
| Maestro | -0.38 | 1.60 |

**Table 4.** Performance on S611 for a subset of 152 mutations with a calculated ΔΔG and with experimental temperatures ranging from 24°C - 26°C and pH ranging from 6.8 - 7.2.

| Method | Pearson Correlation | RMSE (kcal/mol) |
| --- | --- | --- |
| Dynamut2 | 0.79 | 0.95 |
| **PSP_GNM** | **0.73** | **1.07** |
| DUET | 0.69 | 1.25 |
| MCSM | 0.65 | 1.34 |
| SDM | 0.60 | 1.76 |
| Dynamut | 0.51 | 1.41 |
| IMutant | 0.32 | 1.62 |
| MUPro | 0.31 | 1.85 |
| Maestro | -0.45 | 1.69 |
| ENCOM | -0.60 | 1.95 |

## Discussions

The fundamental assumption underlying PSP-GNM is that during unfolding there exists a difference in the order of contacts broken involving the mutation position in the wildtype and mutant protein. For a wildtype-mutant pair, when the contacts are ranked in the order of which they break and only those contacts sharing the same rank are considered, the cumulative difference in the free energies of these contacts between the wildtype and mutant forms correlate with the experimental ΔΔG, as demonstrated in this study. In this context, the free energy takes into consideration the interaction energy between a residue pair given by the Miyazawa-Jernigan statistical potential and the entropy, which is calculated as the mean-squared fluctuation in distance between a residue pair.

In this study, we have used a weighted Gaussian network model that uses Hookean springs of variable stiffness to simulate specific interactions between a residue pair. Such an approach of weighting the Hookean springs between interacting residue pairs based on their identity and chemical nature has been previously shown to improve the ability of elastic network models to capture the near-native protein dynamics [29]. We have demonstrated through PSP-GNM that while maintaining a good agreement with the experimental B-factors, such weighted formulations of elastic network models can also discriminate between the native and mutant forms of a protein. The partial unfolding approach used in our study is a modification of a previous method described by Su et al [20] that was successfully able to predict the order of contact break events. Additionally, to verify the extent of concordance of PSP-GNM with the original GNM we randomly picked two proteins (PDB IDs 1AKY and 1CUN, chain A) from our dataset and obtained the residue-residue cross-correlation heat maps for the wildtype using both methods (Fig. S8). A visual inspection suggests close correspondence between the cross-correlation maps obtained from the two methods.

A major limitation of PSP-GNM is its inability to provide a calculated ΔΔG if no contacts involving the residue at the mutation position are broken in the wildtype and/or in the mutant. To further explain such cases, we randomly selected two wildtype-mutant protein pairs from the S350 dataset and observed that the residue contacts for the mutation positions are primarily within the same secondary structure elements (Fig. S9). Residue-residue contacts within secondary structures are energetically stronger and less easily broken than tertiary contacts, possibly explaining why they were not broken during partial unfolding. As an attempt to address this limitation, we re-defined partial folding by increasing the cutoff to 60%. That is, we considered a protein to partially unfolded if 60% of the native state contacts are broken. However, we observed that the model becomes unstable by generating more than one zero eigen values upon eigen decomposition of the Kirchhoff matrix, suggesting a sparse contact matrix. Consequently, for such cases we assign a theoretical ΔΔG value of 0. Based on the results presented for the S2298 (Fig. 3B), S350 (Fig. 4B) and S611 (Fig. 5B) datasets, we do not observe a considerable reduction in the agreement to experimental ΔΔG when assigning PSP-GNM-calculated ΔΔG value of 0 for such cases.

Three key observations are made in our study. First, we observe variation in the extent of agreement of the calculated ΔΔG with the experimental ΔΔG for different experimental temperatures and pH. We demonstrate that the agreement is strongest for mutants having experimental temperatures close to 25°C and near-neutral pH. Specifically, for the S350 and S611 datasets we observe the correlation is higher for PSP-GNM than for most other methods. Previous studies have also used higher weights for measurements made in the same temperature and pH conditions [3,30]. It is to be noted that PSP-GNM doesn’t take into consideration the experimental temperature to weight the interactions between residues. Based on the observed temperature dependence however, using experimental temperature as a parameter for weighing residue-residue interactions could further improve the extent of agreement with experimental ΔΔG. Second, considering just the contacts broken with the mutation position and their order is key to the reasonably good correlation observed. It strongly suggests of the importance of geometric and energetic changes localized at the mutation site in predicting the overall change in thermodynamic stability of proteins. Finally, a superior performance compared to nine other methods is seen on the dataset of reverse mutations (S611), which is indicative of the ability of PSP-GNM to maintain the anti-symmetric relationship between forward and reverse mutations.

## Conclusion

Understanding the thermodynamic stability changes induced by point mutations in proteins facilitates in understanding the role of mutant proteins in various diseases. Numerous machine-learning-based methods have been developed to predict ΔΔG associated with point mutations. Many of these methods can predict the effect of multiple point mutations. In this study, we have presented PSP-GNM, a novel approach utilizing the knowledge of intrinsic protein dynamics to quantify the thermodynamic changes associated with single amino acid substitutions in proteins. PSP-GNM differs from many of the existing methods in the aspect that it doesn’t use a machine learning approach. Through PSP-GNM we have introduced a formulation of GNM in which interacting residues are weighted by their interaction energy obtained from the Miyazawa-Jernigan statistical potential. PSP-GNM utilizes the knowledge of putative contacts broken during partial protein unfolding to estimate ΔΔG. Although there exist several methods that can predict ΔΔG by simply using the protein sequence, PSP-GNM however relies on protein structure to calculate ΔΔG. Additionally, the present implementation of PSP-GNM can only calculate the ΔΔG for single amino acid changes i.e., for cases where the wildtype and mutant sequences differ by only a single amino acid.

We compared the performance of PSP-GNM with nine existing methods on two different datasets: S350 and S611. On the S350 dataset, PSP-GNM showed a correlation of 0.51 and showed better agreement than five other methods. On the same dataset, for mutants with experimental temperatures close to 25°C and neutral pH, PSP-GNM showed remarkably better performance than six other methods with a correlation 0.65. The superior performance of PSP-GNM on the S611 dataset comprising both forward and reverse mutants is noteworthy. PSP-GNM ranked second with a correlation of 0.58 and 0.73 for all mutations and mutations with experimental temperatures close to 25°C and neutral pH, respectively. Our method also highlights the role of near-native protein dynamics in predicting changes to thermodynamic stability. Importantly, the superior performance of PSP-GNM to other methods incorporating proteins dynamics information (e.g., ENCOM) is suggestive of the importance of including the information on local unfolding and contact changes to predict stability changes upon mutations.

## Data Availability

All datasets used in this study are available at: https://github.com/sambitmishra0628/PSP-GNM

## Disclaimer

The author contributed to this article in his personal capacity. The views expressed in this article are his own and do not necessarily represent the views of the National Cancer Institute.

